# sigfit: flexible Bayesian inference of mutational signatures

**DOI:** 10.1101/372896

**Authors:** Kevin Gori, Adrian Baez-Ortega

**Affiliations:** Transmissible Cancer Group, Department of Veterinary Medicine, University of Cambridge, Madingley Road, Cambridge, CB3 0ES, United Kingdom

**Keywords:** mutational signatures, cancer, Bayesian inference

## Abstract

Mutational signature analysis aims to infer the mutational spectra and relative exposures of processes that contribute mutations to genomes. Different models for signature analysis have been developed, mostly based on non-negative matrix factorisation or non-linear optimisation. Here we present sigfit, an R package for mutational signature analysis that applies Bayesian inference to perform fitting and extraction of signatures from mutation data. We compare the performance of sigfit to prominent existing software, and find that it compares favourably. Moreover, sigfit introduces novel probabilistic models that enable more robust, powerful and versatile fitting and extraction of mutational signatures and broader biological patterns. The package also provides user-friendly visualisation routines and is easily integrable with other bioinformatic packages.

## Background

The study of mutational signatures, the distinctive patterns of somatic alterations left by mutagenic processes on the genomes of cells, has acquired considerable scientific prominence in recent years (1, 2). The identification of mutational signatures and the quantification of their activities in a genome offer valuable insight into the molecular processes that operate on the DNA of cancer and normal cells. These may include endogenous mutagenic processes, such as those arising from deficiencies in DNA repair pathways, and exposures to exogenous carcinogens, such as tobacco smoke (2).

Most analyses of mutational signatures so far have focused on patterns of mutations defined over a categorisation of single-nucleotide variants (SNVs). Specifically, 96 trinucleotide mutation categories are defined for SNVs based on base substitution type (6 categories) and trinucleotide sequence context (i.e. the bases immediately 5′ and 3′ of the mutated base; 16 categories). Base substitution types are collapsed so that only pyrimidine reference bases are considered (for instance, guanine-to-adenine and cytosine-to-thymine mutations represent the same base change on opposite strands, so both can be expressed as cytosine-to-thymine, or C>T.) A vector of mutation counts or mutation probabilities across these 96 categories is conventionally known as a mutational catalogue or spectrum (1, 3).

A considerable number of computational methods for mutational signature analysis have entered the scientific literature over the past five years (4, 5). Perhaps most prominent among these is the Wellcome Sanger Institute’s Mutational Signature Framework (later renamed as SigProfiler) (6), which implements the non-negative matrix factorization (NMF) algorithm that has been repeatedly applied to blind source separation problems in biology and other fields (7). This was the first published method for performing inference of mutational signatures *de novo* (a process also known as signature extraction), and its application resulted in the first ‘gold-standard’ collection of consensus mutational signatures, the COSMIC signature catalogue (http://cancer.sanger.ac.uk/cosmic). A notable alternative to this NMF approach is the probabilistic model introduced by the EMu software (8). This implements an expectation–maximisation (EM) algorithm to perform maximum likelihood estimation using a Poisson mutational model, and is able to account for differences in the opportunity for mutations of each class to occur in the sequence. Later tools have been aimed at estimating the proportions in which a set of predefined mutational signatures are present in a collection of catalogues, a more tractable problem known as signature fitting (4, 5). Notably, signature fitting does not share the considerable sample-size demands of signature extraction methods, and is applicable even to individual mutational catalogues. All of these divergent approaches to mutational signature analysis have been, however, prone to recurrent practical drawbacks, the severest of which include data overfitting, convergence to local optima, elevated computational cost, limited ranges of supported mutation categories, insufficient documentation, poor integration with existing statistical programming frameworks, and lack of publication-ready visualisation routines.

Here we present sigfit, a powerful, versatile and user-friendly software package for the analysis of mutational signatures and broader biological patterns, developed in the R programming language. This package provides both NMF-equivalent and alternative Bayesian probabilistic models for extraction and fitting of signatures, including noise-robust models for the analysis of sparse mutational catalogues, such as those derived from weakly mutated tumours or normal somatic cells, as well as models compatible with continuous data (in contrast to discrete count data). In addition, sigfit introduces novel probabilistic models that enable simultaneous fitting and extraction of signatures, aimed at powering both the deconvolution of known mutational patterns and the discovery of rare or previously undescribed patterns. Moreover, the models in sigfit can be seamlessly applied to mutational profiles with arbitrarily defined mutation categories, enabling the analysis of signatures of short insertions and deletions (indels), dinucleotide variants, large-scale structural variants and copy number variants, among others. The package also offers functions for production of publication-quality graphics of input and output data, as well as an extensive user documentation that includes usage examples and test data sets.

## Implementation

### Overview

Developed as an open-source extension for the popular R programming language and environment, sigfit makes use of the Stan (9) statistical modelling platform, and its interface package rstan (http://mc-stan.org/rstan), to carry out Bayesian inference on probabilistic models of mutational signatures. The package is publicly available on GitHub (http://github.com/kgori/sigfit), and can be installed directly from R using the devtools package (10). A detailed vignette is included in the package to illustrate its use.

### Bayesian probabilistic modelling

#### Multinomial models

Four families of probabilistic signature models are implemented in sigfit. The first of these is statistically equivalent to the mathematical formulation of the NMF approach that was introduced by the Wellcome Sanger Institute’s Mutational Signature Framework (6) (hereafter referred to as WSI-NMF) and has since been adopted by several other tools (4). Such a formulation interprets the observed mutational catalogues as an approximate linear combination of a collection of mutational signatures, with the signature exposures representing the mixing proportions. An equivalent statistical formulation can be defined by modelling the mutational signatures as the probability parameters of independent multinomial distributions, and treating the observed mutational catalogues as draws from a mixture of these distributions, with the signature exposures serving as mixture weights. For a general case with *G* catalogues (genomes), *N* signatures and *K* mutation categories, the data and parameters in the model are as follows.

*M*_*G*×*K*_ is the matrix of observed mutational catalogues,

*S*_*N*×*K*_ is the matrix of mutational signatures,

*E*_*G*×*N*_ is the matrix of signature exposures,

*O*_*G*×*K*_ is the matrix of mutational opportunities,

*m*_*g*_ = [*m*_*g*1_, *m*_*g*2_, …, *m*_*gK*_] (the *g*^th^ row of *M*) contains the observed mutation counts in catalogue *g*,

*s*_*n*_ = [*s*_*n*1_, *s*_*n*2_, …, *s*_*nK*_] (the *n*^th^ row of *S*) contains the mutation probabilities in signature *n*,

*e*_*g*_ = [*e*_*g*1_, *e*_*g*2_, …, *e*_*gN*_] (the *g*^th^ row of *E*) contains the signature exposures in catalogue *g*,

*o*_*g*_ = [*o*_*g*1_, *o*_*g*2_, …, *o*_*gK*_] (the *g*^th^ row of *O*) contains the mutational opportunities for catalogue *g*.

In a signature fitting model, the signatures matrix *S* is treated as fixed data and the exposures matrix *E* is inferred, whereas in a signature extraction model both *S* and *E* are inferred.

The mutational opportunities, *O*, represent the opportunity for each mutation type across the relevant sequence (usually, the entire genome or exome) in each catalogue. For the common case of 96 trinucleotide mutation types, these opportunities correspond to the frequencies of each trinucleotide across each genome or exome. This allows accommodating heterogeneity in sequence composition between samples, such as that induced by copy number alterations. One immediate advantage of incorporating mutational opportunities is that the inferred signatures are not contingent on the sequence composition of the genome under study, and hence are directly applicable to both genomes and exomes, and even across species (11), as long as the relevant mutational opportunities are available. Furthermore, sigfit provides a function (‘convert_signatures’) for adapting existing signatures to different sets of mutational opportunities (for example, transforming genome-relative signatures into exome-relative ones). Mutational opportunities are not a feature of the original WSI-NMF method, and were first introduced by the EMu software (8). In sigfit, the use of opportunities is optional, and their omission results in a formulation statistically equivalent to that of NMF.

To illustrate the NMF-equivalent approach to signature inference implemented in sigfit, we first describe a generative process producing mutations in a set of mutational catalogues (Fig. 1A). In this process, *N* conditionally independent mutational signatures are sampled from a Dirichlet distribution parameterised by *α*. For each of the *G* catalogues, a set of exposures (which function as mixture weights) is sampled from a Dirichlet distribution parameterised by *κ*. For each of the *I* mutations occurring in this catalogue, an indicator value *θ*_*gi*_ is drawn from a categorical distribution parameterised by *e*_*g*_, to specify which of the signatures produces the current mutation. Finally, the category of the mutation is drawn from a categorical distribution parameterised by the indicated signature, 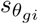. This process is repeated for each mutation in each catalogue.

**Figure 1.**
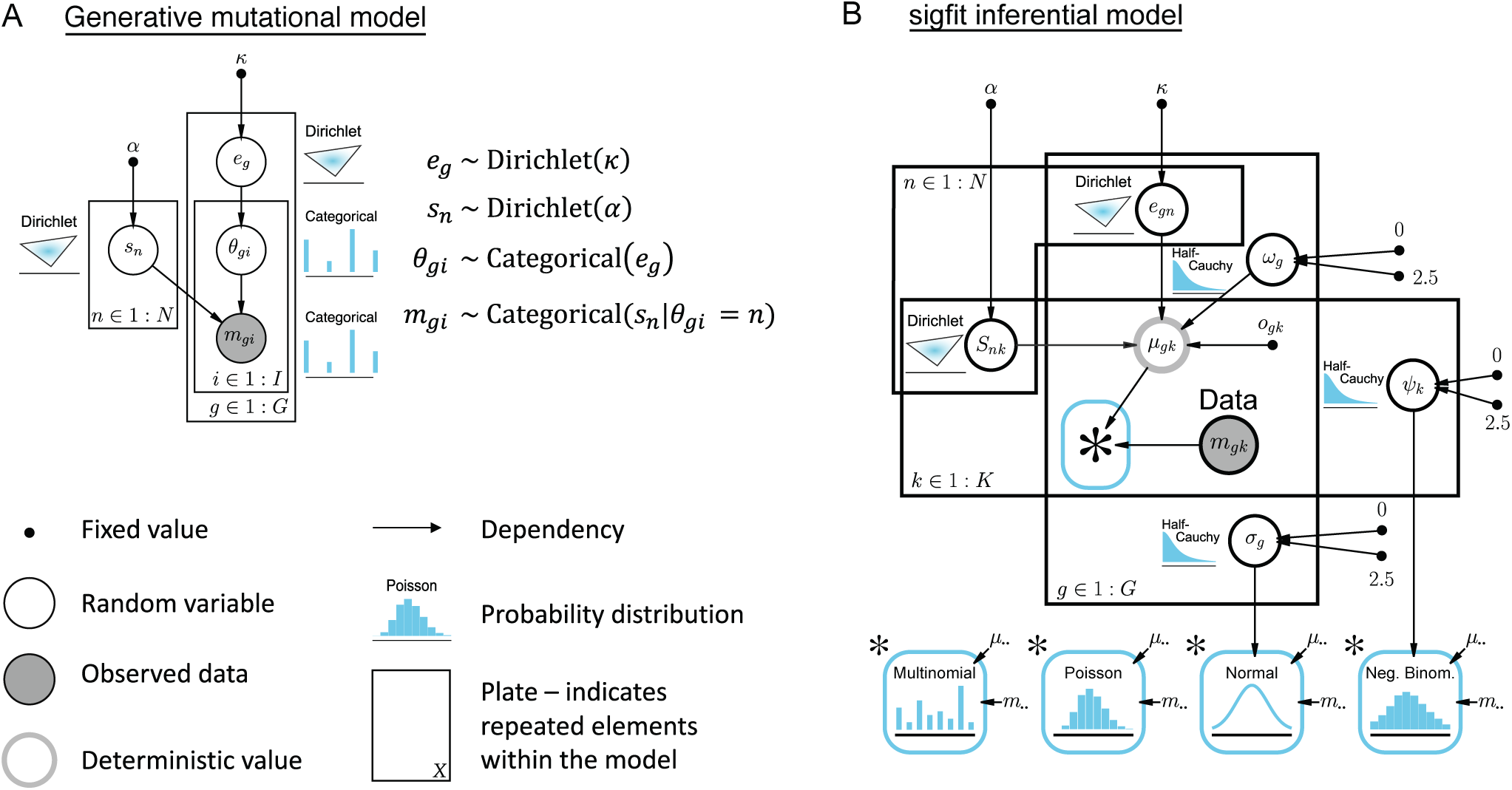
Bayesian model diagrams. **(A)** A generative model of mutation with multiple mutational processes. Each mutational catalogue (indexed by *g*) accumulates *I* mutations. Each mutation *m*_*gi*_ is produced by mutational process *n* of *N* processes. *θ*_*gi*_ encodes the choice of mutational process for mutation *m*_*gi*_ in catalogue *g. θ* is assigned by a draw from a categorical distribution, parameterised by probabilities *e*_*g*_, which are in turn drawn from a Dirichlet distribution parameterised by *κ*. The type of mutation is chosen by drawing from a categorical distribution, parameterised by probabilities *sn*, in turn drawn from a Dirichlet distribution parameterised by *α*. **(B)** The inferential model used by sigfit is based on the generative model, with the distinction that, as the allocation of individual mutations to specific processes is not of interest, the latent parameter *θ* is marginalised out. The process of marginalisation is to matrix-multiply the exposures (*E*, where *eg* is the *g*^th^ row) and the signatures (*S*, where *sn* is the *n*^th^ row), to yield a matrix of expected counts (*μ*, where *μg* is the *g*^th^ row). *μ* is denoted by a node with a grey outline to indicate that it is a deterministic value, distinguishing it from random variable nodes, which take a black outline. The likelihood of the observed mutational catalogue *mg* is obtained from the probability density function of one of the four model classes implemented in sigfit: multinomial, Poisson, normal and negative binomial. Each individual likelihood distribution, and its dependencies, are illustrated below the plate diagram, and can stand in place of the blue box containing an asterisk. Plate notation diagrams were produced using daft-pgm (http://daft-pgm.org).

The inferential model is a simplification of the generative process. Since the assignment of individual mutations to particular signatures is not the aim of mutational signature analysis, considerable computational savings are achieved by marginalising out the assignment and considering only the sufficient statistics (the counts of each mutation category per catalogue), and modelling the likelihood of each catalogue *m*_*g*_ via a multinomial distribution. The parameters of the latter, *μ*_*g*_ = [*μ*_*g*1_,*μ*_*g*2_, …,*μ*_*gK*_], form the *g*^th^ row of a matrix *μ*, which is calculated as the Hadamard (element-wise) product between the mutational opportunities, *O*, and the matrix product of the exposures, *E*, and the mutational signatures, *S* (Fig. 1B*). The model is formalised as follows*.

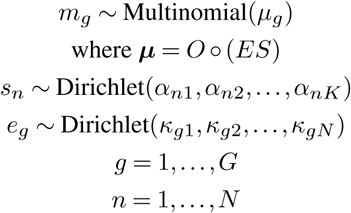

The model provides Dirichlet prior distributions on signatures and exposures via the hyperparameters *α* and *κ*, respectively. By default these are made uninformative by setting all *α*_*nk*_ = 1 and *κ*_*gn*_ = 1. However, users can make use of these parameters to specify custom prior distributions on signatures and exposures.

#### Poisson models

In addition to the multinomial model described above, sigfit offers a number of alternative probabilistic models, one of which is a Bayesian formulation of the Poisson model originally implemented in EMu (8). Here, the mutation counts in each category are modelled as draws from a Poisson distribution with expected value equal to the product of the mutational signatures, the degree of activity of each signature, and the mutational opportunities. For a case with with *G* catalogues, *N* signatures and *K* mutation categories, this model is formalised as follows (Fig. 1B).

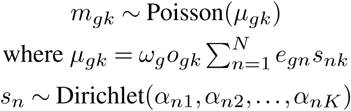

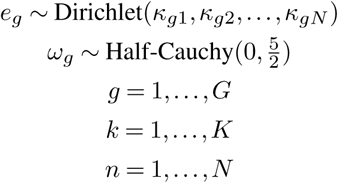

Here, as in the multinomial model above, *m*_*gk*_, *s*_*nk*_, *e*_*gn*_, *o*_*gk*_, *α*_*nk*_, and *κ*_*gn*_ are the elements of matrices of mutations, signatures, exposures, opportunities, and signature and exposure prior hyperparameters, respectively. By contrast, *μ*_*gk*_ represents the expected number (instead of the probability) of mutations of type *k* in catalogue *g*. The terms *ω*_*g*_ serve as catalogue-specific multipliers that allow the transformation of mutation probabilities into expected counts. These multiplier parameters are strictly positive and have weakly-informative half-Cauchy priors.

#### Negative binomial models

The package also implements a noise-robust version of the Poisson model above, in which mutations are modelled using a negative binomial distribution parameterised by an expected mutation count and a mutation type-specific overdispersion, *ϕ* _*k*_. Similarly to the catalogue-specific multipliers *ω* _*g*_, the overdispersion parameters are positive and have vague half-Cauchy priors. For a case with *G* catalogues, *N* signatures and *K* mutation categories, this model is nearly identical to the Poisson model above, except for the data likelihood and the priors for the overdispersion parameters, which are as follows (Fig. 1B).

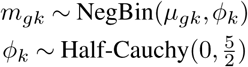

The incorporation of overdispersion provides robustness to sampling noise in the catalogues. This is likely to be beneficial for fitting signatures to sparse catalogues derived from weakly mutated cancers or normal somatic cells, where sampling noise distorts the mutational spectra and prevents optimal fitting of consensus signatures. We note, however, that the incorporation of overdispersion is expected to decrease the computational efficiency of MCMC sampling relative to the Poisson model. Hence, the negative binomial model may be impractical for *de novo* signature extraction in most cases; moreover, sparse catalogues do not normally contain enough information for signature extraction to be feasible.

#### Normal models

The last model family implemented in sigfit comprises signature models based on the normal distribution, with catalogue-specific standard deviations *σ* _*g*_. Similarly to the multipliers *ω*_*g*_, the standard deviation parameters are positive and have vague half-Cauchy priors. For a case with with *G* catalogues, *N* signatures and *K* mutation categories, the formulation of this model is analogous to that of the Poisson model, except for the data likelihood and the priors for the standard deviation parameters, which are as follows (Fig. 1B).

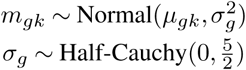

This formulation permits the analysis of signatures from real-valued catalogues, i.e. catalogues containing continuous numeric values instead of discrete mutation counts. Some examples of these are catalogues of methylation beta values obtained from DNA methylation arrays (12), and catalogues representing mutation proportions instead of counts. As for gene expression values, although these are usually expressed on a continuous scale, they tend to be sufficiently large that rounding to the nearest integer entails negligible loss of information. It is therefore preferable to apply the multinomial or Poisson model to this type of data, given the higher efficiency of these models (sigfit automatically rounds input values when a discrete model is selected).

### Combined signature fitting and extraction

In addition to models for conventional extraction and fitting of mutational signatures, sigfit implements a novel probabilistic structure which allows fitting of predefined signatures and extraction of undefined signatures within a single inference process. This formulation, which we term ‘Fit-Ext’, may be understood as a generalisation of the fitting and extraction models described above, where the signatures matrix *S* is composed of *N* signatures known *a priori* (modelled as data), and *M* additional unknown signatures (modelled as parameters). Through multinomial, Poisson, negative binomial and normal formulations analogous to the ones introduced above, the Fit-Ext models perform fitting of the *N* fixed signatures and extraction of the *M* parametric signatures simultaneously. The ‘fixing’ of predefined signatures entails a substantial dimensionality reduction relative to the conventional extraction model, thus enhancing statistical power for deconvolution of any rare or weak signatures that may be present in the mutational catalogues. Fixed signatures may also facilitate the deconvolution of additional signatures that are consistently correlated with the fixed signatures, but which are known to represent distinct mutational processes.

We envision the Fit-Ext models to be of use in three particular scenarios, all of which are exemplified in the Results section below. The first of these scenarios involves the case where a limited sample size precludes signature extraction, yet the data provide qualitative evidence of the presence of one or more novel mutational signatures (not featured in COSMIC or other repositories). By applying a Fit-Ext model to fit the set of known signatures and extract one or more additional signatures from the data, it is possible to infer the novel signatures even from a single mutational catalogue.

The second scenario involves the deconvolution of groups of signatures whose activity is highly correlated across the sample set. A common example of this phenomenon is the case of COSMIC signatures 2 and 13, which correspond to APOBEC-mediated mutagenesis and are almost invariably encountered together. In most sample cohorts, the existence of such correlated groups of signatures precludes their deconvolution, leading to their extraction as a single, composite mutational pattern. Nevertheless, if one of the signatures in the correlated group is fixed using a Fit-Ext model, its accompanying signature or signatures may be effectively deconvoluted, thus increasing the statistical and biological accuracy of the results.

The third scenario for the use of Fit-Ext models involves the identification of rare or weakly present signatures in large cohorts, or in studies of differential mutagenesis (for instance, between primary tumours and their metastases). In these cases, even if signature extraction is applicable, rare signatures may not be amenable to conventional signature extraction due to lack of statistical support. Here, provided that there is evidence of the presence of infrequent mutational patterns which cannot be captured by signature extraction models, a Fit-Ext model may be used to re-fit the previously inferred signatures and extract the rare signatures. Notably, this strategy has recently enabled the deconvolution of a previously undescribed mutational process which operated on the genome of a transmissible cancer in the distant past (13). We note, however, that the approach should be appropriate only when the number of rare signatures is known or suspected; otherwise, the Fit-Ext models are likely to produce signatures composed of uninformative noise.

### Basic usage

sigfit accepts input data as a table of mutations, from which mutational catalogues can be derived using the ‘build_catalogues’ function, or directly as a matrix of catalogues. The mutation table requires at least four fields: sample ID, reference base, mutated (alternate) base, and trinucleotide sequence context of each mutation. If mutations are located in transcribed regions, an additional field can be used to indicate the transcriptional strand of each mutation. We note that the ‘build_catalogues’ function can currently process only mutation data defined over the common 96 trinucleotide mutation types; for data following different categorisations, a matrix of mutational catalogues must be provided.

The main functionality of the package can be accessed via the functions ‘fit_signatures’ (signature fitting), ‘extract_signatures’ (signature extraction) and ‘fit_extract_signatures’ (combined fitting and extraction with Fit-Ext models). These functions carry out Markov chain Monte Carlo (MCMC) sampling on the Bayesian model corresponding to the indicated analysis type and model family. For signature fitting, a matrix of signatures must be provided, whereas for signature extraction, the number of signatures to extract must be specified. If a continuous range of signature numbers (e.g. 2–10) is specified for extraction, sigfit will extract signatures for each value in the range, and suggest the most plausible number of signatures. This is done heuristically, by finding the number of signatures which coincides with the maximum absolute change in gradient of the reconstruction accuracy function (the ‘elbow’ of the curve) (Fig. S1). Reconstruction accuracy is defined, by default, as the cosine similarity between the original catalogues and the catalogues that have been reconstructed using the inferred signatures and exposures.

Recent genomic studies of cancer (14–18) have focused on mutational patterns beyond SNVs, defining mutational signatures over categories of indels, large structural variants and copy number alterations. Whereas some software packages for mutational signature analysis are limited to specific sets of mutation categories (commonly, the 96 SNV categories described above), sigfit can inherently infer signatures over any set of features of interest. Together with the range of available model families and analysis types, this flexibility makes the package suitable for the identification of signatures over a range of unorthodox mutation types, as well as for the analysis of broader biological patterns, or even entirely unrelated signal decomposition problems.

sigfit provides functions to plot mutational catalogues and signatures (‘plot_spectrum’), signature exposures (‘plot_exposures’) and mutational spectrum reconstructions (‘plot_reconstruction’) (Fig. 2A–B). Moreover, all the relevant plots can be produced at once, directly from the output of the signature fitting and extraction functions, using the ‘plot_all’ function. These functions admit not only typical SNV catalogues, but also catalogues with arbitrary numbers of mutation types. A special case is that of transcriptional strand-wise SNV catalogues: if an input mutation table is provided which includes transcriptional strand information, sigfit will automatically define 192 SNV categories, infer strand-wise signatures and adapt its plotting functions correspondingly (Fig. 2C).

**Figure 2.**
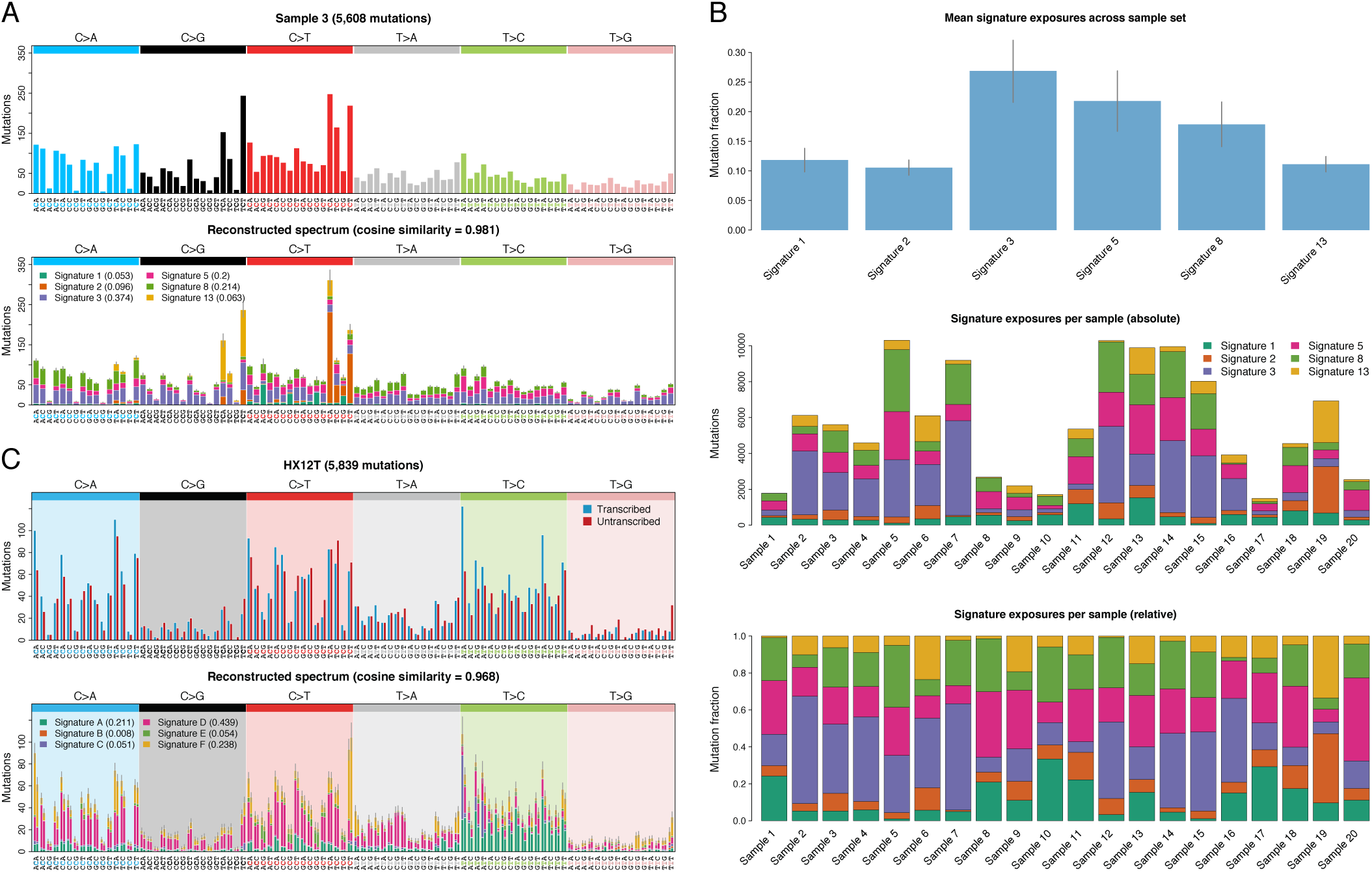
Plotting capabilities of sigfit. **(A)** Plot of the reconstruction of an example mutational catalogue from the estimated signatures and exposures, for a trinucleotide-context SNV catalogue with *K* = 96 mutation types. The original catalogue is shown above the reconstructed one. Colours in the reconstructed catalogue reflect the contributions of each signature. **(B)** Example plots of average signature exposures within a set of catalogues (top), and exposures per catalogue, both as absolute mutation counts (middle) and in relation to the total number of mutations (bottom). **(C)** Plot of the reconstruction of an example mutational catalogue from the estimated signatures and exposures, for a transcriptional strand-wise SNV catalogue with *K* = 192 mutation types. In the original catalogue, shown above the reconstructed one, bars display mutations on the transcribed (blue) and untranscribed (red) strands. All the plots in this figure were produced using the plotting functions in sigfit.

The usage of sigfit is covered in detail in its associated vignette (accessible through the ‘browseVignettes’ function), and in the R function documentation. The package also includes the sets of consensus mutational signatures found in the COSMIC catalogue, as well as several test data sets.

## Results

### Comparison with existing methods

Using previously published mutational catalogues from 119 breast cancer genomes and 88 liver cancer genomes (3), we benchmarked sigfit against the software tools that originally introduced the NMF and Poisson approaches to signature extraction: WSI-NMF (6) and EMu (8).

When comparing the multinomial and Poisson signature extraction models in sigfit to the WSI-NMF and EMu methods, we found that the approaches for estimating the most likely number of signatures in the latter two tools disagreed for both data sets. For the set of 119 breast cancer catalogues, sigfit (both models) and WSI-NMF estimated the most likely number of signatures to be *N* = 6, whereas EMu estimated *N* = 5 as the best value. Conversely, for the set of 88 liver cancer catalogues, sigfit (both models) and EMu estimated the best number of signatures to be *N* = 6, whereas WSI-NMF estimated it as *N* = 10. The contrast arises from the distinct approaches followed by each tool: while sigfit selects the number of signatures maximising the absolute value of the second derivative of the overall cosine similarity between the original and reconstructed catalogues (essentially the ‘elbow method’ used in the clustering literature as a heuristic to estimate the number of clusters in a data set; see example in Fig. S1), WSI-NMF minimises the Frobenius reconstruction error while maximising signature reproducibility (6), and EMu uses the Bayesian information criterion (BIC) to correct for increased model complexity (8). The fact that sigfit consistently estimated the same optimal number of signatures for the multinomial and Poisson models, while inferring highly similar signatures and exposures to those obtained by the other methods (Figs. 3A and S2), suggests that the criterion followed by sigfit to estimate the most plausible number of signatures may be more robust.

**Figure 3.**
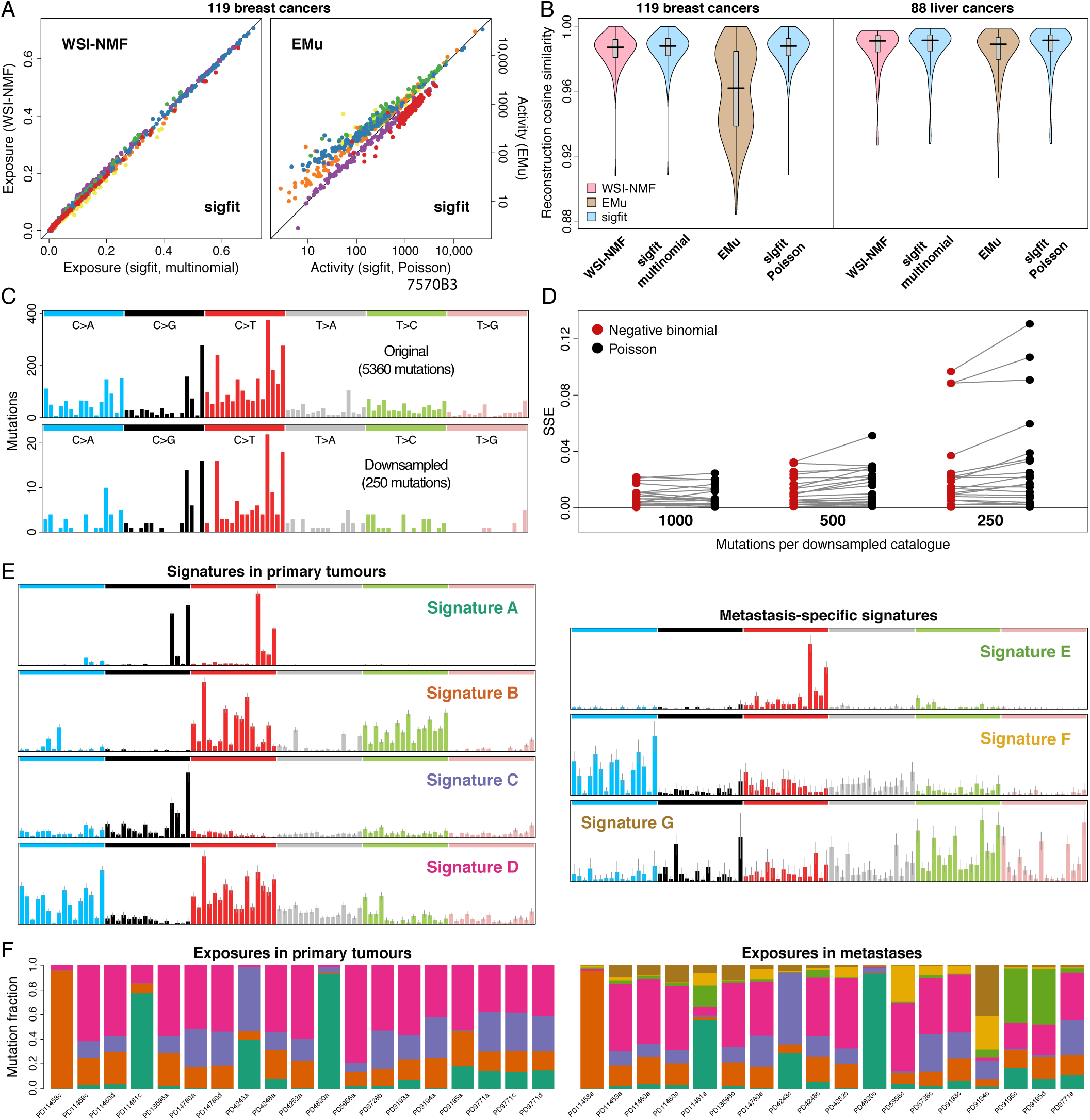
Evaluation of the performance of models in sigfit. **(A)** Left, comparison of signature exposures (proportions of mutations attributed to each signature) as estimated by the multinomial model in sigfit (horizontal axis) and the WSI-NMF software. Right, comparison of signature activities (numbers of mutations attributed to each signature) as estimated by the Poisson model in sigfit (horizontal axis) and the EMu software. The results correspond to extraction of six mutational signatures from a set of 119 breast cancer mutational catalogues (3). Dot colours distinguish between the six mutational signatures. **(B)** Distributions of reconstruction accuracy (cosine similarity between original and reconstructed mutational catalogues) for the models in sigfit and the WSI-NMF and EMu software, in a set of 119 breast cancer catalogues (left) and a set of 88 liver cancer catalogues (right) (3). Boxplots are shown within each violin plot, and horizontal lines indicate median cosine similarity. **(C)** Example of original (top) and downsampled (bottom) mutational catalogues used for testing the performance of the negative binomial model for fitting signatures to sparse catalogues. **(D)** Sum of squared errors (SSE) between the signature exposures inferred by the negative binomial (red) and Poisson (black) signature fitting models when fitting six COSMIC signatures to a set of 21 breast cancer catalogues (19) under different levels of random downsampling. Each dot represents a downsampled catalogue, and lines connect pairs of SSE values for the same catalogue. The number of mutations per catalogue is shown below each group. **(E)** Left, mutational signatures inferred by the multinomial extraction model in sigfit from a set of 19 primary breast tumours (20). Right, signatures inferred by the multinomial Fit-Ext model in sigfit from a set of 18 breast cancer metastases (20), while fitting the four signatures extracted from primary tumours. **(F)** Signature exposures in the primary (left) and metastatic (right) samples used to infer the signatures in (E). Exposure colours match the signature label colours used in (E).

Notably, EMu estimates BIC values and selects the optimal number of signatures prior to signature extraction, whereas both WSI-NMF and sigfit perform extraction for different numbers of signatures and measure the goodness-of-fit of each solution. Together with the fact that the EM algorithm is not guaranteed to converge to a global maximum of the likelihood (21), this can lead to cases where EMu converges to a local maximum for some values of *N*, resulting in incorrect estimation of the number of signatures which best explains the data. This is exemplified by the results for the set of 119 breast cancer catalogues, where EMu converged to a local maximum of the likelihood for *N* = 6 signatures (Fig. S3), hence proposing *N* = 5 as the best number of signatures. The fact that this particular example of convergence to a local optimum was not cherry-picked, but accidentally found, may suggest that this problem is not uncommon. Interestingly, the difference between the solutions reported by sigfit and EMu for *N* = 6 is more prominent in terms of the inferred exposures than the inferred signatures (Figs. 3A and S2). By contrast, WSI-NMF and both models in sigfit found virtually identical solutions for *N* = 6 signatures in this data set (Figs. 3A and S2), which were considerably better than the one reported by EMu in terms of reconstruction accuracy (Fig. 3B). When applied to the set of 88 liver cancer catalogues, however, all the methods showed remarkably similar reconstruction accuracy distributions (Fig. 3B).

It is worth noting that, whereas EMu performs efficient maximum likelihood estimation via EM (8), the WSI-NMF method entails computing a large number of bootstrap replicates in order to identify stable clusters of mutational signatures (6). This causes WSI-NMF to be computationally very expensive, and best suited to highly parallel computing infrastructures. In contrast, the models in sigfit, which exploit the efficient No-U-Turn sampler algorithm for MCMC sampling implemented in the Stan framework (9), incur only moderate memory and CPU demands that are easily met by laptop or desktop computers, while providing virtually identical solutions to those obtained by WSI-NMF (Figs. 3A–B and S2).

In addition to its favourable performance in signature extraction problems, when fitting the entire set of 30 COSMIC signatures to a previously published set of catalogues from 21 breast cancer genomes (19), sigfit reported only six significantly active signatures (Fig. S4), five of which have been previously reported in breast cancer (3, 14). By contrast, optimisation-based methods such as least-squares regression and quadratic programming are notoriously prone to overfitting (22), especially when applied to non-convex functions like those involved in signature extraction and fitting problems. We believe that sigfit’s capability to estimate realistic contributions of known signatures to sets of catalogues (or even single catalogues), without the need to prevent overfitting by constraining the set of candidate signatures, makes the package valuable for small genomic studies that lack the statistical power required for signature extraction.

### Robust signature fitting to sparse catalogues

The noise-robust probabilistic models incorporated in sigfit, which adopt the overdispersed negative binomial distribution to accommodate excess variance in the observed mutation counts, can be used to improve the accuracy of signature fitting to highly sparse mutational catalogues, such as those derived from weakly mutated genomes or exomes. To demonstrate the performance of the negative binomial fitting model, we constructed catalogues with decreasing numbers of mutations by performing multinomial downsampling on the set of 21 breast cancer genomes introduced in the previous subsection (19). This is achieved by sampling catalogues of mutations from multinomial distributions centred on the mutation frequencies observed in the original data. To further increase variance, we added moderate random noise by perturbing each observed mutation frequency value by a random amount *ε*, with |*ε* | < 10^−3^. We obtained three sets of downsampled mutational catalogues, respectively containing 1000, 500 and 250 mutations per catalogue (see example in Fig. 3C). A set of six COSMIC signatures previously reported in breast cancer (signatures 1, 2, 3, 5, 8 and 13) were fitted to each set of catalogues using the Poisson and negative binomial models in sigfit, and the signature exposures estimated by each model for the downsampled catalogues were compared with the exposures estimated by the same model for the original catalogues, by measuring the sum of squared errors (SSE) between both. Whereas, overall, the SSE increases with decreasing number of mutations for both models, the error incurred by the Poisson model grows more rapidly, with the difference between both models being most pronounced for the set of catalogues with 250 mutations (Fig. 3D). In general, the negative binomial model estimated more precise exposures for most of the catalogues in each set: 81% and 57% of catalogues with 500 and 250 mutations, respectively, presented more accurate exposures under the negative binomial model, with median SSE reductions of 33% and 34% relative to the Poisson model, respectively. Moreover, error reductions were greater in those catalogues displaying the largest errors (Fig. 3D). This provides an illustration of the potential of models based on robust probability distributions for the analysis of signatures in sparse and noisy data sets.

### Analysis of metastatic samples using Fit-Ext models

As discussed above, the Fit-Ext models for combined fitting and extraction may be applicable to data sets characterised by qualitative evidence of the presence of rare or novel signatures, the deconvolution of which is precluded by lack of statistical power. This has been demonstrated in our recent analysis of mutational processes in an ancient transmissible cancer, where this methodology was applied to deconvolute the signature of a previously undescribed mutational process which operated on this cancer’s genome in the distant past (13). Here, we present an additional case which may benefit from the application of Fit-Ext models, namely the comparison of mutational catalogues from metastatic cancer samples and their primary tumours. Specifically, to demonstrate the utility of the Fit-Ext models for the identification of subtle mutational patterns, we analysed a previously published set of 19 primary breast tumours and 18 metastases found in the same patients (20).

As shown in the original study of these samples, the mutational processes involved in primary tumours and metastases within each patient are very similar (20). As a result, conventional models for signature extraction infer virtually identical sets of signatures from the 19 primary tumours, the 18 metastases, or the combination of both (Fig. S5). In contrast, using the multinomial Fit-Ext model in sigfit to extract additional signatures from the metastatic catalogues, while fitting the four fixed signatures extracted from the primary tumours, identified mutational patterns that are specific to the metastatic samples (Fig. 3E). In this case, the metastasis-specific signatures do not reflect different mutational processes from those operating in the primary tumours, but rather represent mutational components that are enriched in the metastases relative to the primary tumours, revealing that the four signatures obtained by the extraction model are composite. Specifically, signature E corresponds to COSMIC signature 2, which, despite being represented (together with COSMIC signature 13) by signature A in the primary tumours, shows increased activity in three metastatic samples, two of which belong to the same patient (Fig. 3F). Signature F most likely corresponds to COSMIC signature 8, which is represented by signature D in the primary tumours; exposure to signature F, which is especially prominent in two of the metastases (Fig. 3F), seems to reflect an increased contribution of C>A mutations from signature 8, relative to the pattern in signature D. Finally, signature G, which is only highly active in one metastatic sample (Fig. 3F), might represent a mixture of the T>N components of COSMIC signature 9 (which are also split between signatures B and D) and an unknown C>G component. This analysis demonstrates the Fit-Ext models’ potential for identifying rare mutational patterns which are commonly undetectable to signature extraction approaches, and for revealing latent variability in the mutational components of inferred signatures.

### Further applications of Fit-Ext models

Besides the discovery of rare or novel signatures, the Fit-Ext strategy can also assist the deconvolution of highly correlated pairs or groups of signatures, which are consistently observed to co-occur but are known to reflect distinct mutational processes. Notable examples of this are the pairs of COSMIC signatures 1 and 5, and 2 and 13, which were deconvoluted into individual signatures only through the analysis of thousands of tumours (3). For small sample sizes, current signature extraction methods are often unable to separate these correlated signature pairs (19). The Fit-Ext models in sigfit can assist in the deconvolution process by defining one of the signatures in the relevant pair as fixed, with the effect that its companion signature can be extracted separately. To demonstrate this, we first performed conventional signature extraction using the multinomial model on the same set of 21 breast cancer catalogues used above (19). This produced an optimal set of four signatures, two of which are respective mixtures of COSMIC signatures 1 and 5, and signatures 2 and 13 (Fig. S6A, first and second rows). Then, we applied the multinomial Fit-Ext model to fit one fixed signature, chosen to be COSMIC signature 13, and extract three signatures. The inferred signatures demonstrate how fixation of signature 13 prompts a clean extraction of signature 2 (Fig. S6B). Finally, signatures 1 and 5 were separated by defining the former as a second fixed signature, and hence inferring two signatures highly similar to COSMIC signatures 2 and 5 (Fig. S6C). This illustrates the Fit-Ext models’ ability to prompt the deconvolution of groups of known, highly correlated signatures, leading to more-precise signature exposure estimates and greater biological accuracy of the inferred signatures. The approach may prove particularly useful in studies involving small or highly homogeneous collections of catalogues.

In addition, we assessed the effectiveness of the Fit-Ext approach in cases where signature extraction is not possible, yet there is evidence for the presence of a novel mutational pattern in the data. To this end, we repeatedly applied the multinomial Fit-Ext model to individual simulated mutational catalogues, which were drawn from a multinomial distribution defined as a random mixture of three COSMIC signatures (signatures 1, 2 and 7) and one simulated novel signature (Fig. S7A). The novel signature was drawn from a Dirichlet distribution with low concentration values (*α*_*k*_ = 0.05, *k* = 1, …, 96), and bore no resemblance to any signature in COSMIC (all cosine similarities < 0.327). In each of 100 simulation experiments, sigfit was applied to the artificial catalogue in two ways. First, the multinomial signature fitting model was used to fit all 30 COSMIC signatures to the simulated catalogue; this resulted in inaccurate catalogue reconstruction (median cosine similarity between original and reconstructed catalogues, 0.887), as none of the COSMIC signatures reflected the features of the novel signature (Fig. S7B). Second, the multinomial Fit-Ext model was applied to fit all COSMIC signatures to the simulated catalogue, while extracting one additional signature. Because a single catalogue was used in each simulation, the original and reconstructed catalogues were always identical, as the reconstruction error is captured by the extracted signature (Fig. S7B). Despite this, the simulated novel signature was accurately extracted in the vast majority of cases (median cosine similarity 0.948; Fig. S7C). This illustrates the utility of Fit-Ext models in scenarios where signature fitting cannot completely capture the mutational patterns in the data.

### Analysis of methylation patterns using normal models

To provide an example of the application of signature models based on the normal distribution to the analysis of real-valued biological data, we examined DNA methylation data from 50 breast cancer samples and 27 solid normal tissue samples, retrieved from the TCGA-BRCA project at the NIH National Cancer Institute’s Genomic Data Commons Data Portal (http://portal.gdc.cancer.gov). We selected open-access samples for which methylation beta values derived from the Illumina Human Methylation 27 array were available; the set of tumours was randomly subset to 50 samples (Table S1). For these 77 samples, we retained only data for probes corresponding to CpG sites in the promoters of genes which were labelled in the COSMIC Cancer Gene Census (http://cancer.sanger.ac.uk/census) as having a germline or somatic association with breast cancer. In addition, we ignored probes with variance < 10^−4^. The resulting 99 probes were adopted as mutation categories, and mutational catalogues were defined as vectors of beta values over such probes in each sample. Mutational signatures were extracted from these catalogues using the normal extraction model in sigfit, for *N* = 1, …, 10 signatures, using default priors and no mutational opportunities. The most informative results were obtained for *N* = 7 signatures (Fig. 4A). It should be noted that these patterns do not constitute *bona fide* mutational signatures, as individual CpG sites cannot be considered to be actual mutation types. Instead, the inferred signatures represent raw patterns of gene promoter methylation; although they do not directly reflect differential methylation between cancer and normal genomes, differential methylation patterns can be assessed by examination of the estimated signature exposures (Fig. 4B). Even in this moderately sized set of samples, methylation differences are such as to enable clustering of tumours and normal tissues into distinct groups (with the exception of one normal sample; Fig. 4B). Remarkably, tumours exhibit substantially higher heterogeneity in promoter methylation than normal samples, with variable hypermethylation of well-studied cancer genes including *BRCA1, BRCA2, CASP8, CDH1, ESR1, GATA3* and *NTRK3*, and hypomethylation of oncogene *CCND1* (Fig. 4*). In addition, the extracted signatures suggest concurrent methylation of multiple genes in some tumours, involving genes such as BRCA1, BRCA2, NTRK3, PBRM1* and *RB1* (‘Sig. A’ in Fig. 4A).

**Figure 4.**
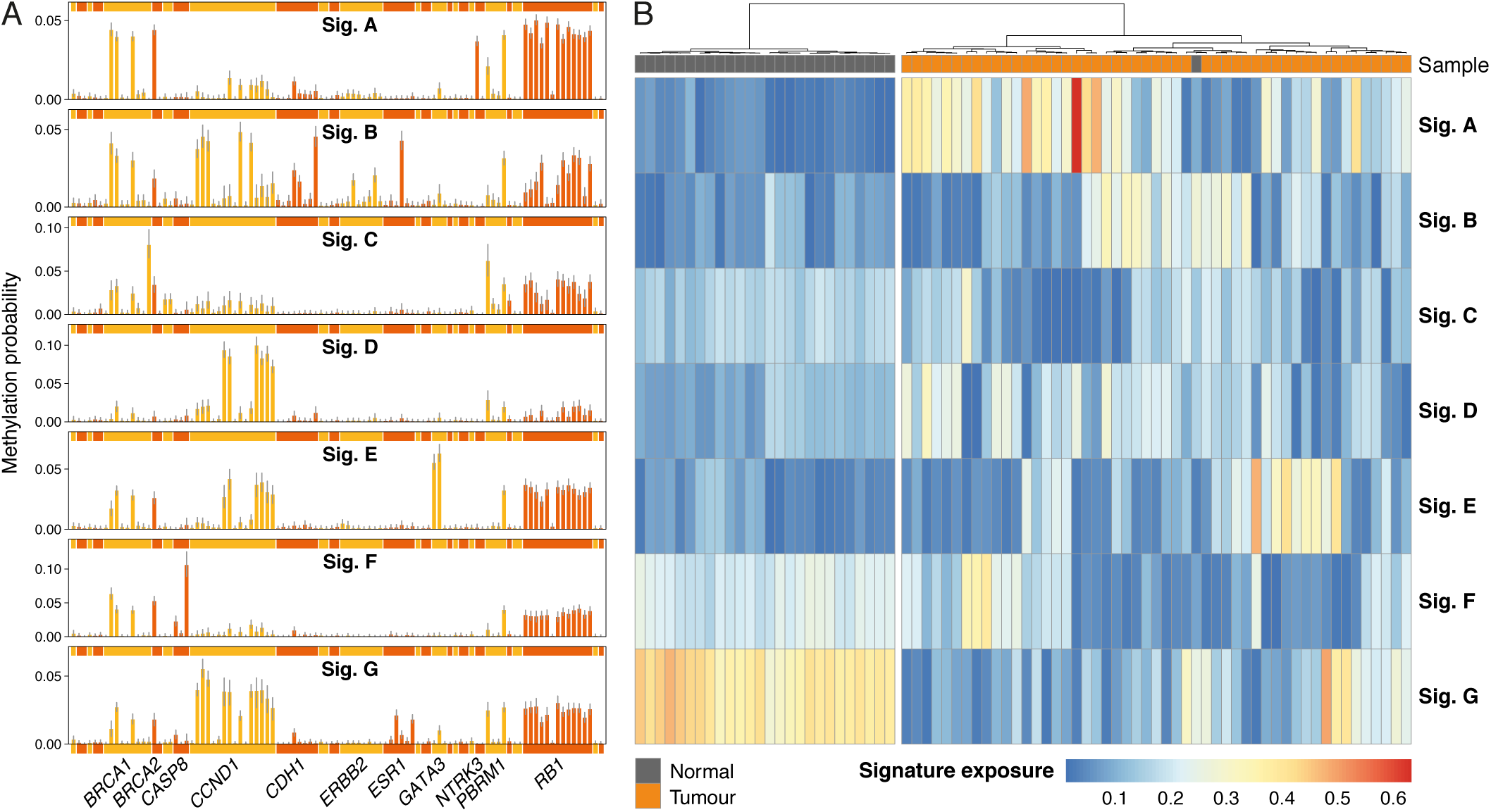
Patterns of gene promoter methylation in tumour and solid normal tissue samples from breast cancer patients. **(A)** Seven signatures inferred using the normal extraction model in sigfit (labelled Sigs. A–G), representing DNA methylation patterns extracted from catalogues of methylation beta values at 99 CpG sites located in the promoter regions of breast cancer-associated genes. Each bar presents the methylation probability of a CpG site; alternating yellow and orange colours are used to distinguish groups of sites in the same gene. Selected gene symbols are shown along the horizontal axis. **(B)** Heatmap of inferred exposures to signatures A–G in 27 normal tissue samples and 50 breast cancer samples (indicated by grey and orange rectangles along the upper margin, respectively). Unsupervised sample clustering was performed using Ward’s method on the Euclidean distance between signature exposures. The heatmap was produced using the pheatmap R package (23).

It is important to note that the present analysis does not constitute sufficient evidence of the potential biological role of any of the mentioned promoter methylation events in breast cancer. Instead, the results above are intended as an illustration of the effectiveness of versatile probabilistic signature models for the analysis of biological patterns beyond established mutational signatures, including those to be found in continuous data.

## Discussion

Although computational approaches for mutational signature analysis have proliferated rapidly, many existing methods exhibit recurrent conceptual or practical limitations, including excessive rigidity of supported data formats (e.g. many tools admit only SNV data), constrained analysis types (e.g. few tools offer both signature fitting and extraction), excessive computational requirements, convergence to local optima, and poor user-friendliness. By exploiting the Bayesian inference machinery offered by the Stan statistical framework (9), the sigfit package is designed to provide easy, flexible and robust analysis of mutational signatures and other biological patterns. We have demonstrated the comparability of the probabilistic models implemented in sigfit to the most prominent methods for mutational signature analysis, and have illustrated the versatility and variety of analyses made possible by the new methodologies introduced by the package. Examples of practical value include improved accuracy of signature fitting to sparse mutational catalogues using noise-robust models, enhanced deconvolution of known and novel signatures by means of combined fitting and extraction, and analysis of patterns in continuous biological data, such as those arising from DNA methylation assays. We have also illustrated the flexibility and ease-of-use of the publication-quality visualisation routines included in the package. Moreover, the usefulness of the functionalities in sigfit has been recently evidenced by its adoption by other research groups (24–26).

We believe that the novel methodologies implemented in sigfit constitute a significant step towards simpler and more adaptable analysis of mutational signatures and broader biological patterns. Furthermore, these functionalities are underpinned by the popularity and portability of the R programming environment, which greatly facilitate reproducibility and integration of data and results with other bioinformatic analysis workflows.

## Software availability

sigfit is publicly available on GitHub (http://github.com/kgori/sigfit), and is licenced under the GNU General Public License v3.0.

## Acknowledgements

The authors would like to thank Maximilian Stammnitz for testing the software, and Daniel Gaffney for helpful suggestions and bug reports.

---

**Figure S1.**
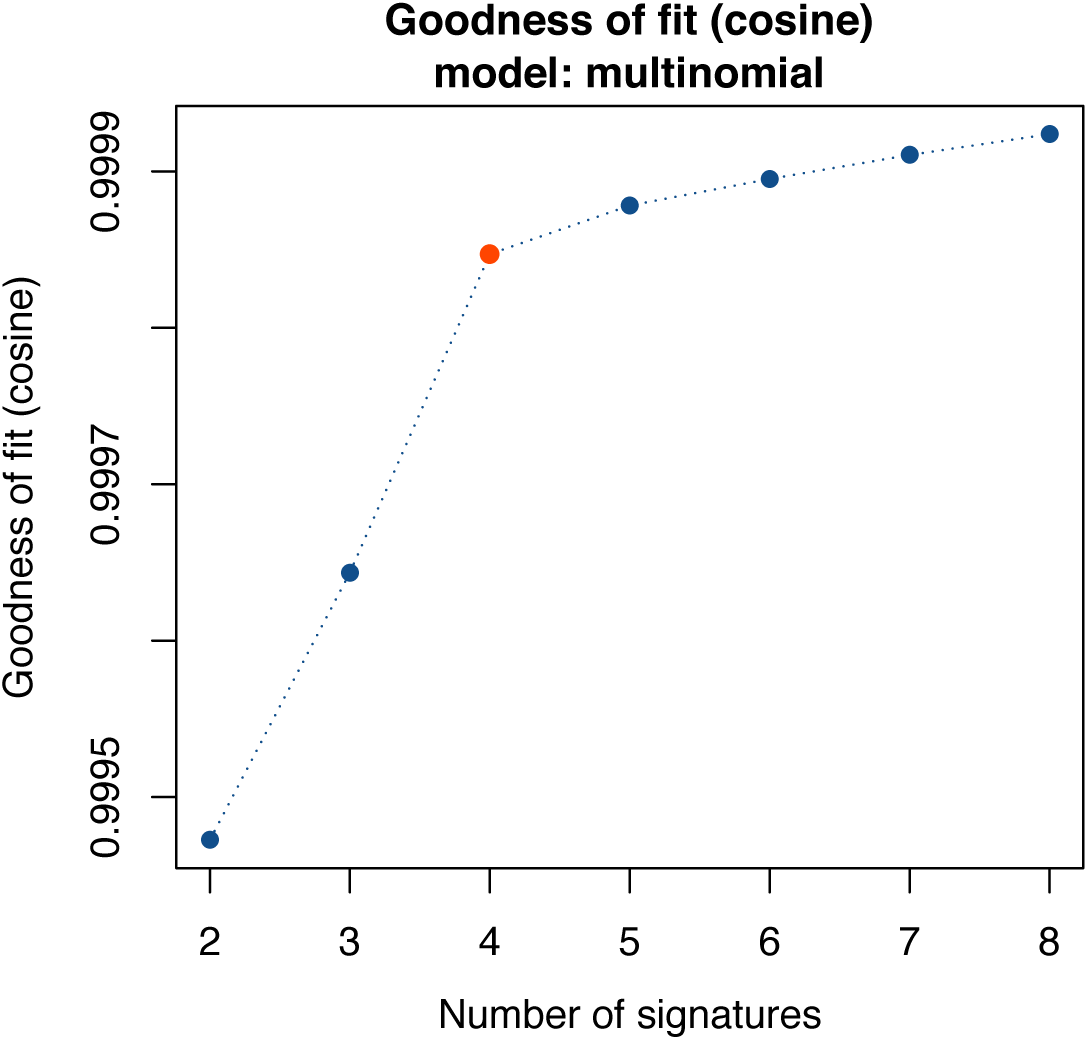
Signature number selection via goodness-of-fit. Graph displaying spectrum reconstruction accuracy (measured as the cosine similarity between the original catalogues and those reconstructed from inferred signatures and exposures) for a range of numbers of signatures, as produced by the ‘plot_gof’ function in sigfit. In this example, a value of four signatures is automatically suggested as the point maximising the approximate second derivative of the curve (indicated in orange).

**Figure S2.**
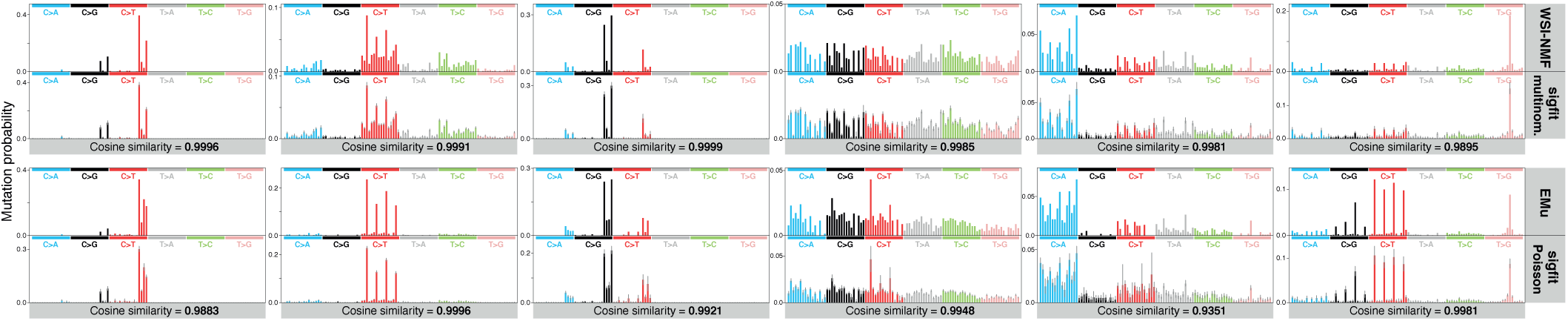
Mutational signatures extracted by sigfit and existing methods. Comparison of mutational signatures extracted *de novo* by WSI-NMF and the multinomial extraction model in sigfit (top row), and by EMu and the Poisson extraction model in sigfit (bottom row) from a set of 119 breast cancer catalogues (3). Cosine similarity between analogous pairs of signatures is shown below each pair. Vertical axes present mutation probabilities, and bars represent 96 trinucleotide mutation types, with colours indicating base substitution type.

**Figure S3.**
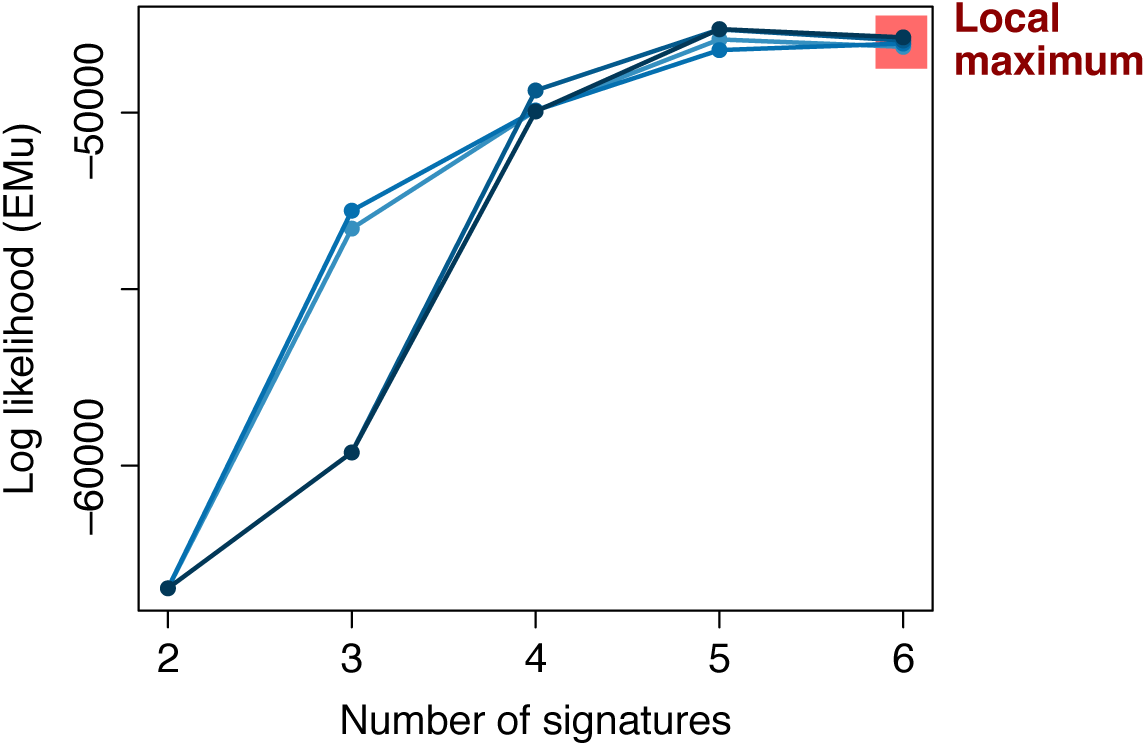
Convergence of EMu to local maximum of the likelihood. Graph displaying the log-likelihood calculated by EMu over a range of numbers of signatures, for five different runs of the software (indicated by lines in different colours, two of which overlap). The decrease in likelihood when extracting six signatures (highlighted in red), compared to five signatures, indicates convergence of the expectation–maximisation algorithm to a local maximum of the likelihood.

**Figure S4.**
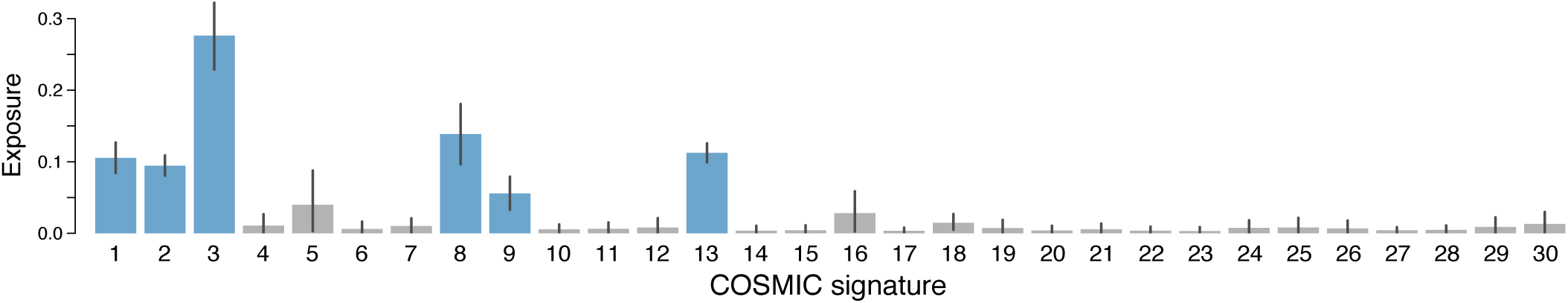
Accurate fitting of consensus mutational signatures to cancer catalogues. Posterior mean estimates of global signature exposures inferred by the multinomial signature fitting model in sigfit, through fitting of the 30 signatures in the COSMIC catalogue to a set of 21 breast cancer catalogues (19). Error bars indicate 95% highest posterior density (HPD) intervals. Grey bars indicate non-significant signature exposures, defined as exposures for which the lower limit of the 95% HPD interval is below 0.01.

**Figure S5.**
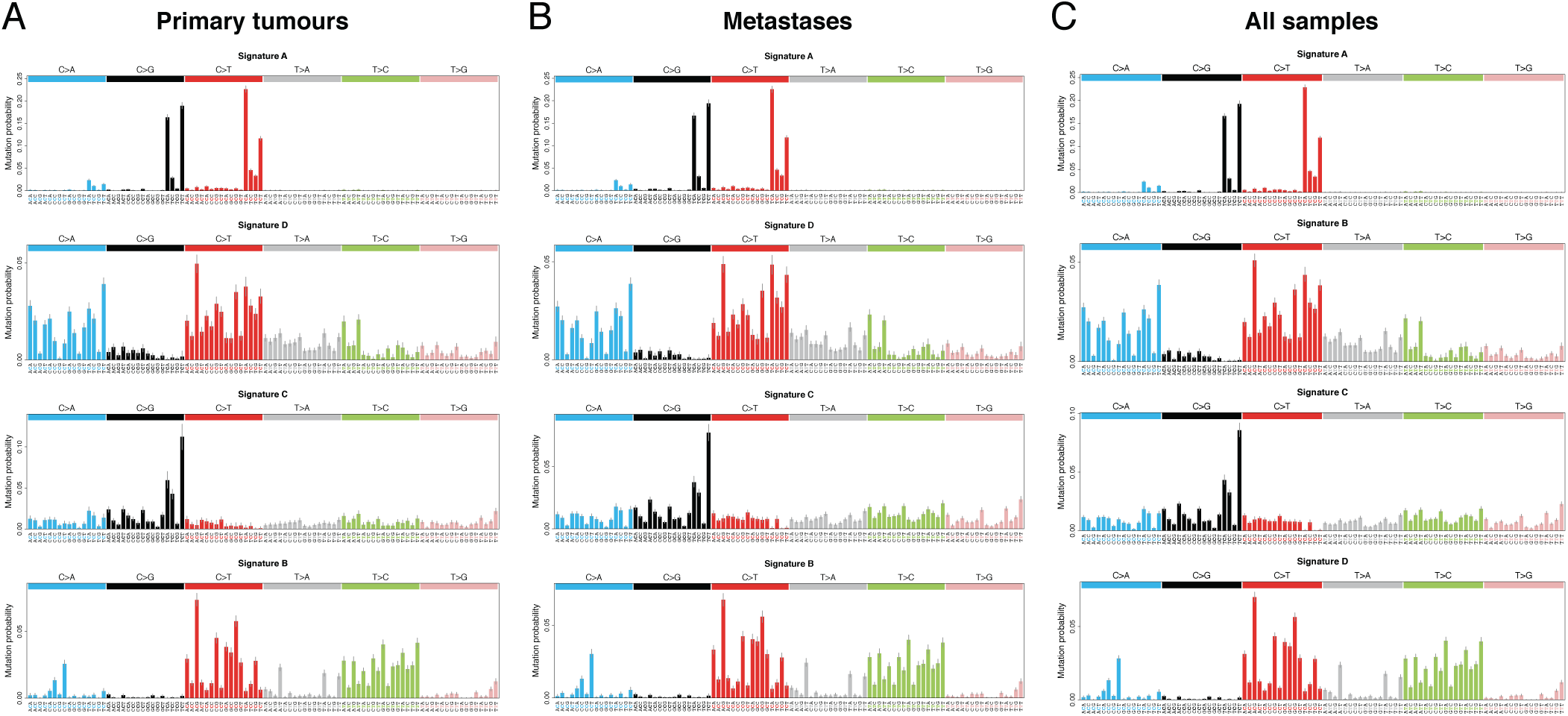
Mutational signatures extracted from primary breast tumours and metastases. Sets of four signatures inferred from primary and metastatic breast cancer samples (20) by the multinomial signature extraction model in sigfit. Signatures were extracted from **(A)** 19 primary breast tumours, **(B)** 18 metastatic samples obtained from the same patients, and **(C)** primary and metastatic samples combined. In all cases, sigfit suggested four as the most plausible number of signatures.

**Figure S6.**
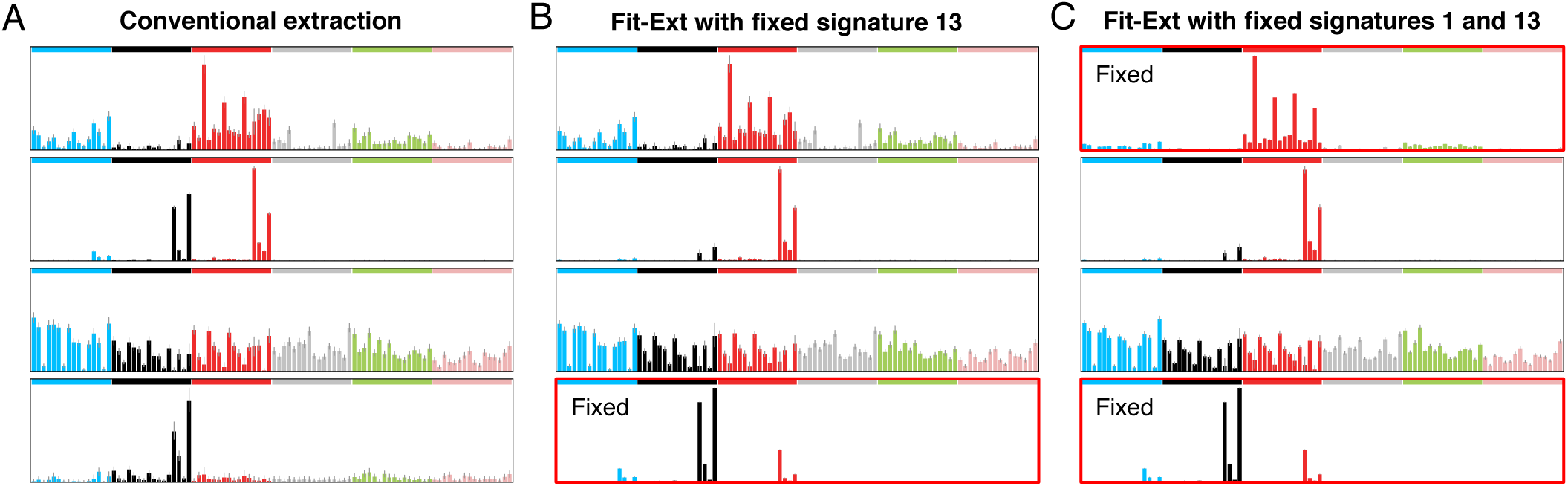
Deconvolution of correlated signatures using Fit-Ext models. Sets of four mutational signatures extracted by sigfit from 21 breast cancer catalogues (19) using **(A)** the multinomial extraction model, **(B)** the multinomial Fit-Ext model with fixed (pre-specified) COSMIC signature 13, and **(C)** the multinomial Fit-Ext model with fixed COSMIC signatures 1 and 13. Fixed signatures are highlighted in red.

**Figure S7.**
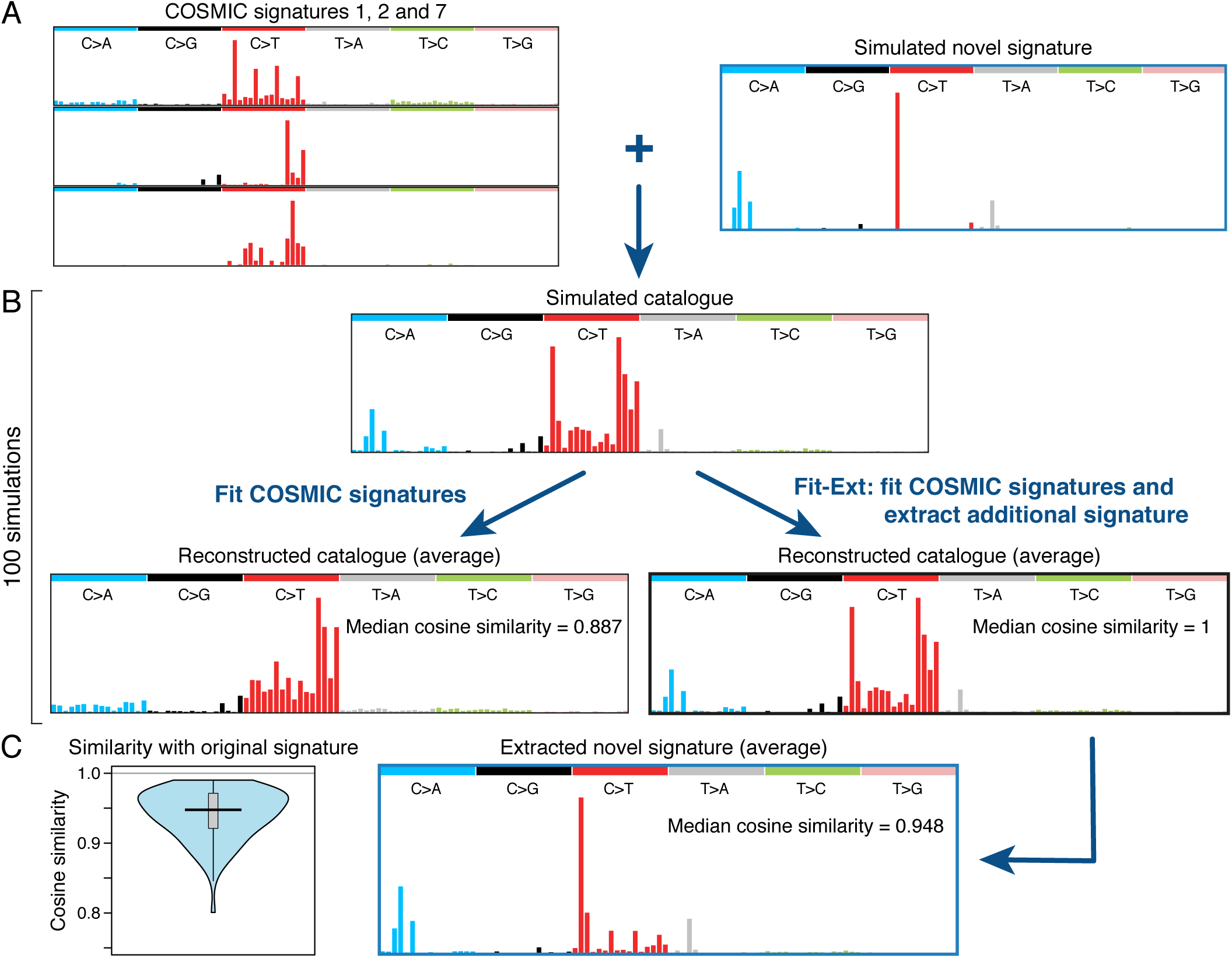
Evaluation of the Fit-Ext model on simulated mutation data. **(A)** Mutational spectra of the three COSMIC signatures (left) and the simulated novel mutational signature (right) that were used to produce simulated mutational catalogues. **(B)** In each of 100 simulations, a mutational catalogue (top) was generated from a random mixture of the signatures shown in (A), and signature analysis was performed with sigfit via two separate approaches: using the multinomial fitting model to fit all COSMIC signatures to the catalogue (bottom left), and using the multinomial Fit-Ext model to fit all COSMIC signatures and extract one additional signature (bottom right). Average reconstructed mutational catalogues (constructed from the product of signatures and inferred exposures) and median cosine similarities between the original and reconstructed catalogues across all simulations, are shown for each approach. **(C)** Average mutational spectrum of the estimated novel signature, as inferred by the Fit-Ext model across all simulations. The violin plot (left) presents the distribution of the cosine similarity between the original novel signature, shown in (A), and the inferred signature across all simulations. A boxplot is shown within the violin plot, and the horizontal line indicates the median cosine similarity.

